# *pyfraglib*: An integrated cfDNA fragmentomics platform

**DOI:** 10.64898/2026.07.22.740152

**Authors:** Daniel Schuette, Laura K. Godfrey, Jessica Schneider, Sven Borchmann, Jan-Michel Heger, Roland F. Schwarz

## Abstract

**Summary:** Cell-free DNA (cfDNA) fragmentomics is the analysis of a diverse set of cfDNA fragment features, e.g. fragment length profiles, windowed protection scores, and end motifs. As such it requires software tooling for fragment extraction, statistical feature modeling, and cohort-level comparative analysis. *In silico* simulations can facilitate the development and validation of new methods by generating testing datasets with known ground truth. Existing tools address individual aspects of this workflow but none provide all necessary capabilities within a single package.

**Results:** We present *pyfraglib*, a platform integrating fragment extraction from short- and long-read sequencing, statistical feature modeling (Gaussian mixture and NMF decomposition of fragment length profiles, end motif diversity, windowed protection scores), cohort-level differential testing of said features, and a simulation module. The library is exposed through a command-line interface, a Python API, and a Nextflow pipeline. We demonstrate *pyfraglib* in two ways. First, on two simulated 20-sample cohorts we show that pyfraglib’s per-sample and cohort-level analyses recover the differences introduced by construction. Second, we apply *pyfraglib* to 88 cfDNA samples from a central nervous system lymphoma (CNSL) study and construct a fragmentomics score combining an NMF signature with end motif and WPS summaries via a classifier trained on cerebrospinal fluid and healthy donor plasma samples. As a proof of concept and applied to 66 baseline patient plasma samples, the score identifies a high-risk subgroup with worse failure-free survival (log-rank p=0.0247).

**Conclusions:** *pyfraglib* integrates sample- and cohort-level fragmentomics analyses as well as *in silico* simulation within a consistently engineered Python framework. *pyfraglib* source code and documentation are available at https://github.com/schwarzlab-ccb/pyfraglib.

## Background

The analysis of cell-free DNA (cfDNA), obtained through liquid biopsy, offers minimally invasive insights into physiological and pathological states such as pregnancy and cancer [1, 2]. More specifically, the non-random fragmentation of cfDNA results from the interplay of multiple biological factors including nuclease activities, chromatin structure, and cell death mechanisms [1]. Fragmentation patterns differ systematically between healthy and diseased states, between tissues of origin, and across different pathological processes [3, 4]. These characteristics collectively constitute the fragmentome and include but are not limited to fragment length distributions, 5′ end motifs, and patterns of genomic coverage coinciding with nucleosome occupancy and gene expression [4, 5].

Several studies have demonstrated the utility of fragmentomics features for tumor detection and cancer progression monitoring, particularly when multiple features are combined via machine learning [6]. Importantly, fragmentomics can be an orthogonal source of information in scenarios where mutation-based risk assessment is infeasible. This includes tumors with low mutation burden as well as early-stage tumors where circulating tumor DNA (ctDNA) levels are comparatively low relative to the overall pool of cfDNA [7–9].

As cfDNA fragmentomics analyses have received increasing attention, computational tools (Table 1) have been developed to facilitate automatic feature extraction from sequencing data, often with a specific focus on certain types of features or specific data modalities. For example, frameworks such as cfDNAPro [10] and FinaleToolkit [11] support the computation of multiple fragmentomic features, including fragment length distributions, end motif frequencies, and genome-wide protection scores. In contrast, Griffin [12] is a dedicated tool for nucleosome profiling based on windowed protection scores with GC-bias correction, FrEIA [13] focuses on fragment-end sequence diversity for cancer detection, and LBFextract [14] extracts entropy-based transcription factor activity signals.

**Table 1:**
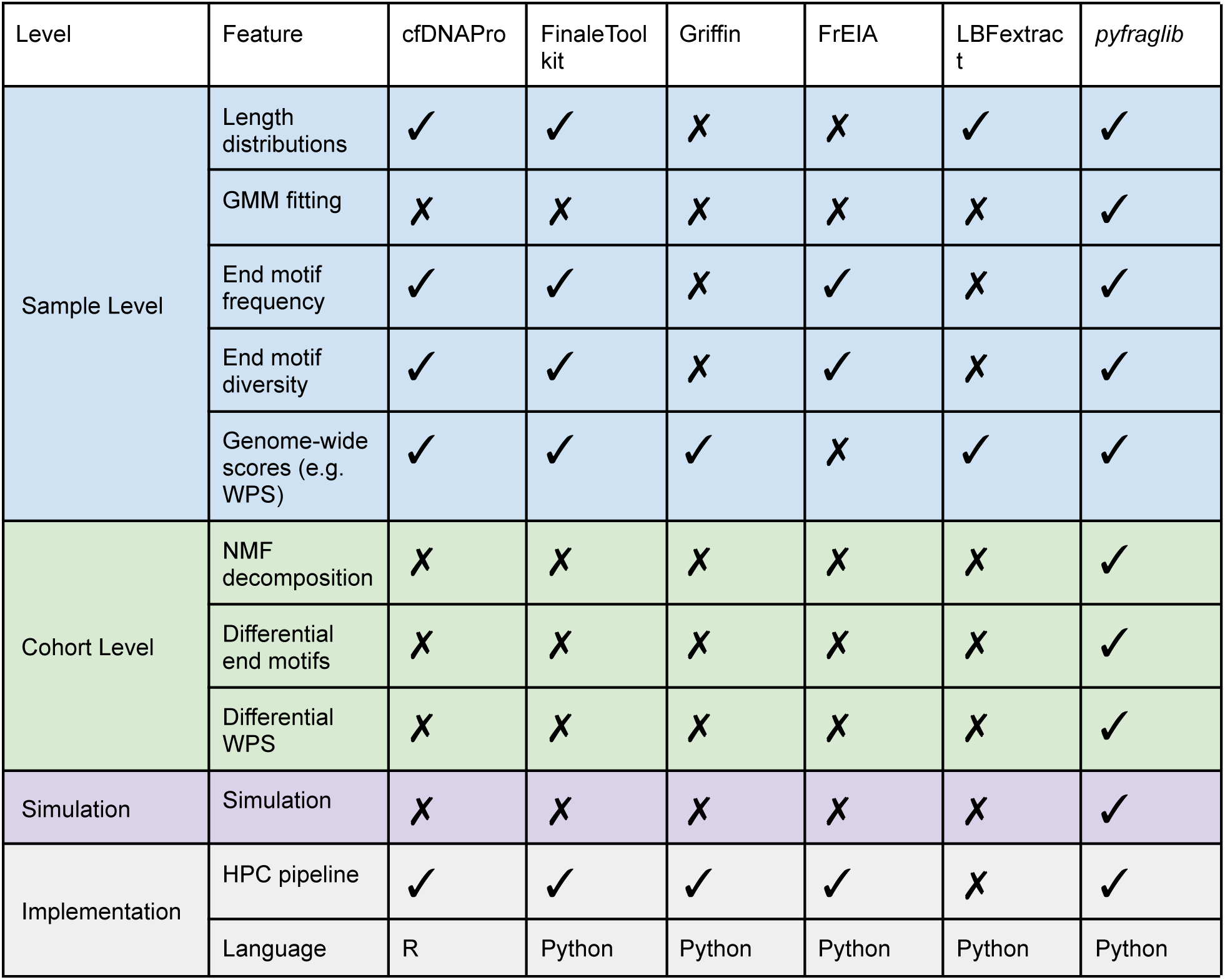
Method comparison table of fragmentomics tools. ✓: supported; ✗: not supported or not reported in the cited publication.

While these tools have substantially simplified extraction of cfDNA fragmentomics features, their specialised nature and limited support for cohort-level analyses is a challenge to their routine application in clinical cohorts, where reproducibility is key. In particular, integrated methods for identifying differential fragmentomic features between groups, decomposing cohort-wide fragmentation signatures, and generating interpretable summary statistics suitable for downstream prognostic or predictive modeling are largely absent. As a result, cohort-level analyses are frequently implemented through bespoke, study-specific scripts, limiting reproducibility and software reuse.

A second, related gap is the absence of *in silico* frameworks for generating synthetic cfDNA with biologically informed nuclease, tissue, and chromatin models. Such infrastructure is needed because certain cohort-level methods like NMF are unsupervised, and their behaviour on a given mixture of cfDNA features cannot be validated against ground truth from patient samples alone. While simulation is a mature calibration tool in adjacent areas of computational genomics [15], comparable infrastructure is missing for cfDNA fragmentomics.

We address these gaps with *pyfraglib*, a Python library that combines, in a single consistently engineered package: (i) a sample-level feature-extraction module exposed through a Python API, a JSON-configured CLI, and a Nextflow HPC pipeline; (ii) cohort-level inference modules for calculating Gaussian mixture and NMF decompositions of length profiles as well as differential end-motif and WPS testing; and (iii) a biology-informed cfDNA simulator that parameterises nuclease activity, tissue mixture, chromatin state, and fragment-length distributions to provide the controlled-ground-truth setting needed to calibrate cohort-level methods during development. We first simulate two cohorts of 20 samples each to illustrate how *pyfraglib*’s simulation module can aid in method prototyping. Then, we analyze 88 cfDNA samples from a central nervous system lymphoma cohort [16] to show how an NMF signature as well as end-motif and WPS scores can be combined into a cerebrospinal fluid (CSF)-likeness score that identifies a high-risk subgroup with worse failure-free survival.

## Implementation

### Overview of *pyfraglib*

*pyfraglib* implements an end-to-end workflow for analyzing fragmentomics features both from real-world or simulated sequencing data. *pyfraglib* is organized into three main modules: (i) a sample-level feature-extraction module, (ii) a cohort-level inference module and, (iii) a basic biology-informed cfDNA simulation module for prototyping and benchmarking (Figure 1). As input data to the sample-level feature-extraction module, users provide aligned BAM files from either short- or long-read sequencing, and optionally VCF files containing known single nucleotide variants. Alternatively, simulated data generated by the simulation module can be used as input source.

**Figure 1:**
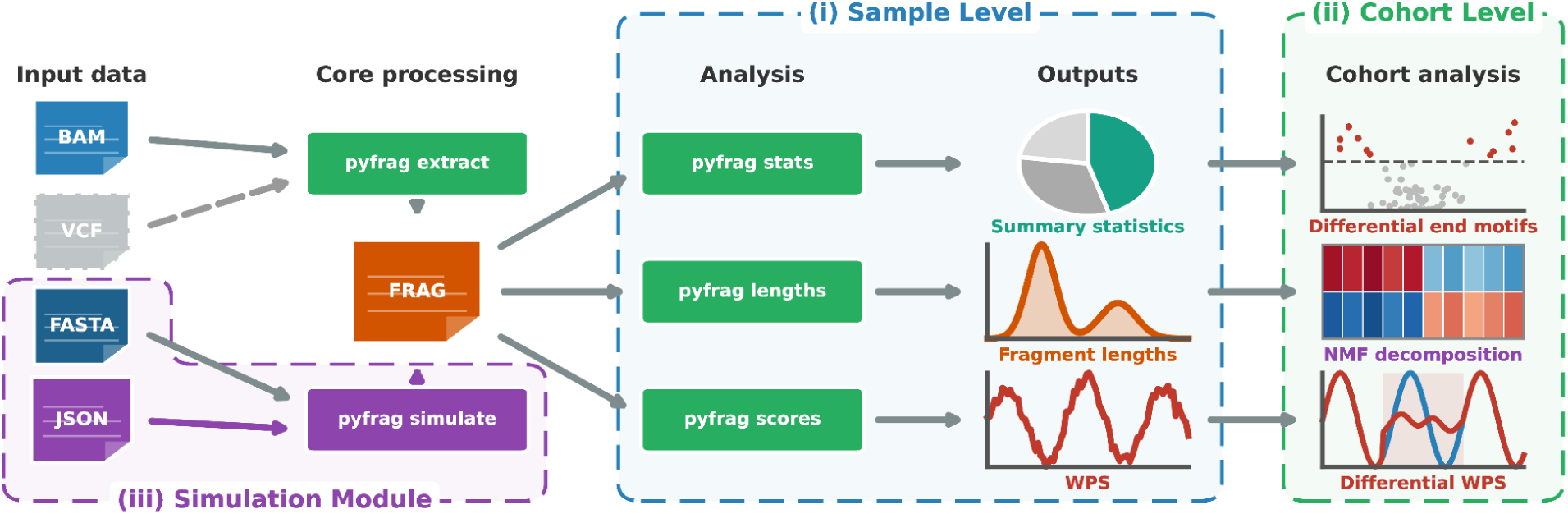
pyfraglib workflow overview. The pyfraglib command line interface supports two input sources: real sequencing data (BAM file, with optional variant calls in VCF format) are processed by the **extract** subcommand into the compact binary FRAG file format, while synthetic data are generated by the **simulate** subcommand from a reference genome (FASTA) and a JSON configuration file. All downstream analyses operate on FRAG files through three main subcommands: **stats** computes per-sample summary statistics like fragments per chromosome, **lengths** fits Gaussian mixture models to fragment length distributions, and **scores** calculates genome-wide fragmentomics scores like the windowed protection score (WPS) and motif diversity scores. Additional commands that come with pyfraglib allow for cohort-level analyses, more specifically differential end motif assessment, NMF decomposition of fragment length profiles, and differential analysis of WPS.

The application programming interface (API) is organized into three primary layers: fragment processing, feature extraction and analysis, and simulation (Supp. Figure 1a). The command line interface (CLI) subcommands follow these layer naming conventions and deal with individual samples. Cohort-wide analyses are implemented in stand-alone Python scripts. Analyses and simulation are configured through JSON files for reproducibility and for large-scale analyses, a Nextflow pipeline (Supp. Figure 1b) automatically parallelizes per-sample fragment extraction and feature calculation, then aggregates results for cohort-level analyses.

### Sample-level feature-extraction module

*pyfraglib’*s sample-level feature-extraction module performs the extraction of fragment information (length, end motifs and optionally mutation status) from single- or paired-end BAM files from long- or short-read cfDNA sequencing with or without a user-provided VCF file. It then stores the results in a FRAG file, a novel, dedicated and highly compressed binary file format for efficient storage and retrieval (see *Data storage and compression* and Supp. Fig. 1). These FRAG files serve as the common input for the three implemented feature analyses: fragment length profiling, end-motif analysis, and windowed protection score (WPS) analysis. For each analysis, *pyfraglib* computes the corresponding feature summaries and generates diagnostic plots, which are saved in the output together with the processed data. All resulting outputs can be used directly as input to the cohort-level inference module, avoiding repeated parsing of the original alignment files and enabling efficient, asynchronous downstream analyses.

The fragment length distribution of cfDNA exhibits a characteristic multimodal pattern, reflecting the underlying nucleosomal organization of DNA wrapped around nucleosomes and the mechanisms of DNA fragmentation during cell death [1, 4]. The dominant peak typically corresponds to mono-nucleosomal fragments (∼167 bp), while additional enrichment around multiples of this length represents di-nucleosomal (∼334 bp) and higher-order nucleosomal fragments. To model these fragment length profiles, *pyfraglib* implements a Gaussian mixture model (GMM)-based approach, where the observed fragment lengths are represented as a weighted combination of multiple Gaussian components. The model is fitted using the expectation-maximization (EM) algorithm [17], which iteratively estimates component parameters by alternating between assignment of fragment lengths to latent Gaussian populations (expectation step) and optimization of component means, variances, and mixture proportions (maximization step). Component counts, initialization bounds and model complexity can be configured through JSON files provided to the **lengths** subcommand, giving users explicit control over the fitted model. An example configuration is provided in Table 2. The fitted GMM captures mono-nucleosomal (∼167 bp), di-nucleosomal (∼334 bp), and additional fragment populations, with component means, standard deviations, and mixing weights saved to JSON for downstream analysis.

**Table 2:**
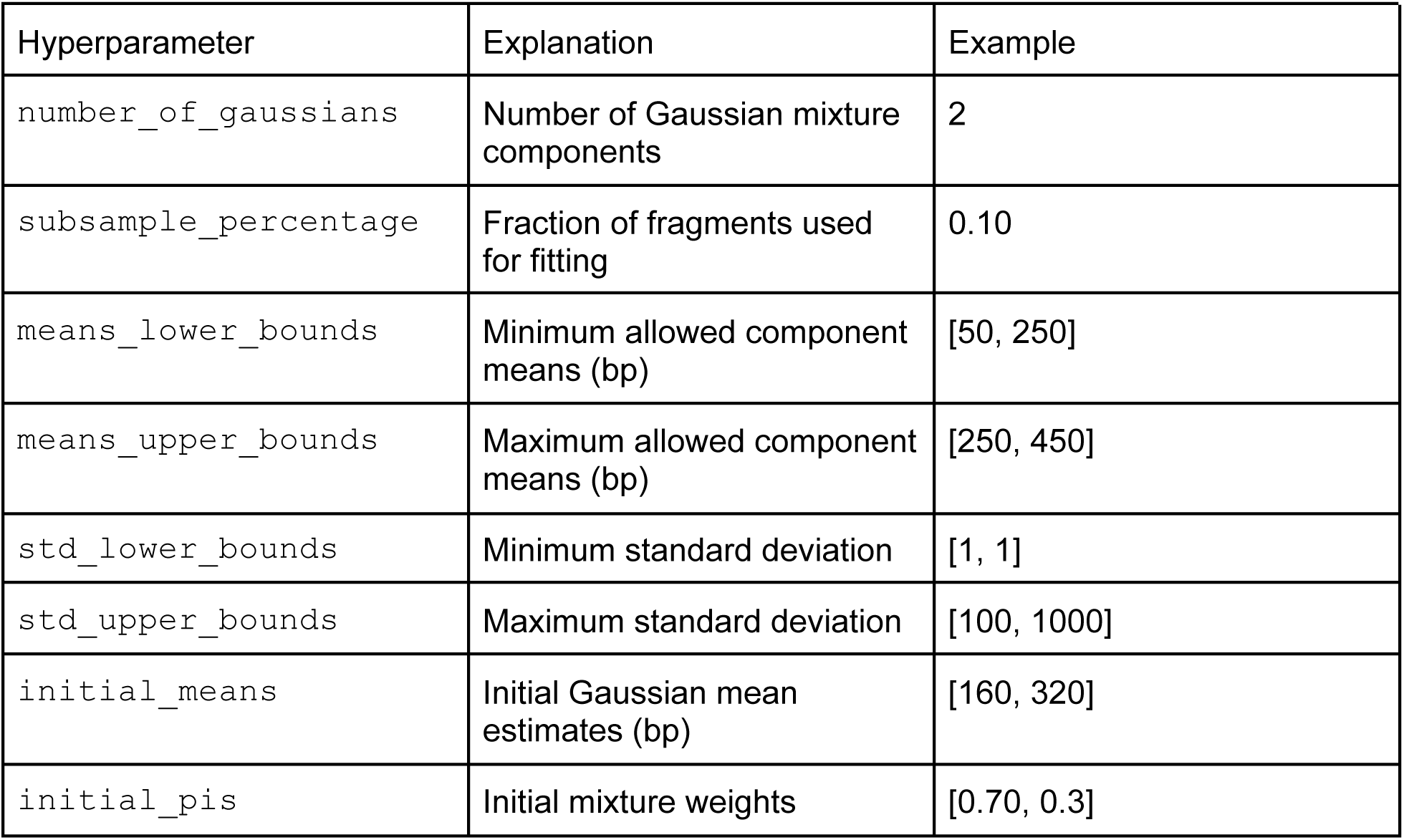
Example JSON configuration file for the GMM fitting of fragment length profiles using a 10% subsampling and two components.

| Hyperparameter | Explanation | Example |
| --- | --- | --- |
| `number_of_gaussians` | Number of Gaussian mixture components | 2 |
| `subsample_percentage` | Fraction of fragments used for fitting | 0.10 |
| `means_lower_bounds` | Minimum allowed component means (bp) | [50, 250] |
| `means_upper_bounds` | Maximum allowed component means (bp) | [250, 450] |
| `std_lower_bounds` | Minimum standard deviation | [1, 1] |
| `std_upper_bounds` | Maximum standard deviation | [100, 1000] |
| `initial_means` | Initial Gaussian mean estimates (bp) | [160, 320] |
| `initial_pis` | Initial mixture weights | [0.70, 0.3] |

In addition to fragment length distributions, cfDNA molecules exhibit non-random sequence features at their fragment ends, which reflect the enzymatic and structural processes involved in chromatin fragmentation [1, 4]. To characterize these terminal sequence patterns, *pyfraglib* provides the **stats** subcommand, which extracts k-mer-based 5′ and 3′ end motif frequencies from cfDNA fragments. To summarize the overall diversity and complexity of end-motif profiles within each sample, *pyfraglib* calculates complementary diversity metrics, including Shannon entropy [18], which measures the uncertainty and distributional complexity of observed motifs, and the Simpson index [19], which emphasizes the dominance of highly abundant motifs.

Genome architecture strongly influences cfDNA fragmentation by the physical protection of DNA wrapped around nucleosomes from enzymatic degradation, leading to characteristic depletion and enrichment patterns of fragments originating from nucleosome-occupied regions [4]. To quantify nucleosome-associated protection patterns, *pyfraglib* implements the windowed protection score (WPS) algorithm [4] in the **scores** subcommand. Users specify genomic regions of interest via BED files to limit computational scope. For each genomic position in the BED file regions, WPS is calculated within a sliding 120 bp window as the difference between the number of fragments that fully span the window and the number of fragments with at least one fragment endpoint occurring within the window. Regions protected by nucleosomes are therefore expected to exhibit increased WPS values due to the depletion of fragment ends within protected intervals, whereas accessible or nucleosome-depleted regions show reduced protection scores. This approach converts fragment-level positional information into a continuous genomic signal describing local chromatin protection patterns.

The resulting raw per-position WPS signal is post-processed by *pyfraglib* to reduce technical noise. First, rolling-median subtraction is applied using a default 1001 bp window to remove long-wavelength drift, caused by regional coverage variation and other low-frequency effects. Second, a Savitzky–Golay filter (21 bp window, polynomial order 3) [20] is applied to suppress high-frequency Poisson sampling noise while preserving the local shape and periodic structure of nucleosome-associated oscillations.*To* summarize the strength of nucleosomal phasing within each analyzed region, *pyfraglib* transforms the normalized WPS signal into the frequency domain using the discrete Fourier transform [21]. The resulting power spectrum is used to calculate the ratio of the mean spectral magnitude in a narrow band around 1/170 cycles per bp to the mean magnitude across all frequencies. Under white noise, this ratio has expectation 1, while values above 1 indicate signal concentration at the nucleosomal phasing wavelength and therefore stronger evidence of ordered nucleosome-associated fragmentation. Treated as a scalar feature, this ratio enables quantitative comparison of chromatin organization patterns between cfDNA samples.

### Cohort-level inference module

The cohort-level inference module enables integrative analysis of fragmentomic features across multiple samples using outputs generated during feature extraction in the feature-extraction module. All three supported feature classes - fragment length distributions, end-motif profiles, and windowed protection scores - can be jointly analyzed at the cohort level to identify shared fragmentation patterns and statistically significant differences between sample groups. In addition to the per-sample feature files, the module requires a cohort annotation table containing sample identifiers together with the corresponding BAM, BAI, and, optionally, VCF file locations.

To identify latent fragmentation patterns within a cohort, *pyfraglib* applies non-negative matrix factorization (NMF) [22] to the sample-by-feature fragment length matrix. NMF factorizes the non-negative input matrix V into two lower-dimensional non-negative matrices, W and H, such that V≈WH. Here, W contains the contribution (or exposure) of each latent fragment length signature in every sample, while H represents the characteristic fragment length profile associated with each signature. Previous studies have demonstrated that these latent signatures capture biologically meaningful cfDNA fragmentation patterns that can distinguish physiological and pathological states [23]. The inferred signature and exposure matrices are exported together with diagnostic visualizations to facilitate downstream interpretation and quality assessment.

Cohort-level analysis of end motifs is performed using non-parametric differential testing to identify sequence motifs whose abundances differ significantly between predefined sample groups. For each observed k-mer, *pyfraglib* compares motif frequencies using the Wilcoxon rank-sum test [24], followed by Benjamini–Hochberg false discovery rate (FDR) correction [25] to account for multiple hypothesis testing. The resulting statistics provide effect estimates and adjusted *P*-values for each end motif, enabling systematic identification of group-specific fragmentation patterns.

Differential analysis of chromatin organization is performed analogously using regional WPS-derived summary statistics. For each genomic region, *the power spectrum ratio outlined above* is compared between sample groups using the Wilcoxon rank-sum test with Benjamini–Hochberg FDR correction. Regions are defined using the annotation field provided in the BED file supplied during WPS computation within the feature extraction module. When multiple genomic intervals share the same annotation, their corresponding WPS-derived measurements are aggregated prior to statistical testing, enabling region-level comparisons across biologically meaningful genomic features rather than individual intervals.

### cfDNA simulation module

In addition to analyzing experimental sequencing data, *pyfraglib* provides an integrated simulation module for generating synthetic cfDNA fragment datasets with user-defined statistical properties enabling method development, benchmarking, algorithm validation, and controlled evaluation of fragmentomic analyses. The generated output follows the same FRAG file specification as experimental data and can therefore be processed by the sample-level feature extraction and cohort-level inference modules without modification.

Simulation is based on a user-specified reference genome sequence in FASTA format together with a JSON configuration file defining the statistical model parameters (Supp. Figure 1c). Fragment lengths are sampled from a configurable Gaussian mixture distribution and end motifs are generated by probabilistically cutting along the reference sequence. Cleavage probabilities are determined by a parametric nuclease model describing the sequence preferences and relative activities of the three principal nucleases implicated in cfDNA fragmentation, namely DNASE1L3, DNASE1, and DFFB [26]. The model therefore captures both the characteristic fragment length distribution and the sequence-dependent end-motif composition of cfDNA molecules.

To model heterogeneous biological samples, the simulator additionally supports mixtures of two independently parameterized fragmentation processes. Rather than using a single set of simulation parameters, users may specify two complete parameter sets together with a mixing fraction controlling the relative contribution of each population to the final dataset. This enables simulation of biologically relevant scenarios, such as circulating tumor DNA (ctDNA) diluted within a background of healthy cfDNA. Because each component is generated independently before mixing, both fragment length distributions and end-motif profiles can differ between the constituent populations. The parameters used to simulate the two cohorts in this study are listed in Table 3. The Supplementary Documentation describes the probabilistic fragment length and end motif generating processes in greater detail.

**Table 3:** Simulation hyperparameters for the two synthetic cohorts. Cohort 1 consists entirely of background fragments. Cohort 2 is an 85%/15% mixture of background and tumor-like fragments. SD = standard deviation.

| Hyperparameter | Background | Tumor-like |
| --- | --- | --- |
| DNASE1L3 activity | 8.0 | 0.5 |
| DNASE1 activity | 2.0 | 2.5 |
| DFFB activity | 0.2 | 5.0 |
| Mononucleosomal mean (bp) | 167 | 150 |
| Mononucleosomal SD (bp) | 10 | 18 |
| Mononucleosomal fraction | 0.85 | 0.92 |
| Dinucleosomal mean (bp) | 334 | 300 |
| Dinucleosomal SD (bp) | 30 | 30 |

### Data Storage and compression

*Pyfraglib’s* sample-level feature extraction module extracts per-sample fragments and writes them to disk as FRAG files. Thereby, *pyfraglib* achieves a substantial reduction in file size by using the binary file format Apache Parquet (https://parquet.apache.org/, o. J.). FRAG files only retain information required for fragmentomics analysis, which is fragment coordinates, lengths, and end motifs, while discarding the underlying sequencing reads. This design not only improves storage efficiency and I/O performance but also reduces the amount of potentially identifying genetic information contained in the files. Nevertheless, FRAG files are not fully anonymized. Although they do not contain complete DNA sequences, the combination of genomic fragment coordinates and end-motif information could, in principle, still permit donor re-identification. Consequently, FRAG files should be handled with the same care as other human genomic data and shared in accordance with applicable ethical and regulatory guidelines.

### Availability, Reproducibility, and Testing

*pyfraglib* is distributed under the GNU General Public License v3, with source code available at https://github.com/schwarzlab-ccb/pyfraglib. The codebase requires Python 3.12+ and enforces strict type annotations (PEP 484) with static type checking through mypy. Code style is maintained PEP8 compliant through flake8 linting. Correctness is verified by unit and integration tests, covering all core functionalities. A Nextflow [27] pipeline enables parallel distributed processing across HPC clusters with SLURM or LFS scheduling. Documentation is generated through Sphinx from inline docstrings, ensuring consistency between code and documentation.

## Results

We demonstrate *pyfraglib*’s capabilities in two complementary ways. First, we simulated two synthetic cohorts and then used *pyfraglib*’s per-sample and cohort-level analyses to recover the prescribed differences against this known ground truth (Figures 2 and 3). Second, we applied *pyfraglib* to 88 cfDNA samples from a central nervous system lymphoma (CNSL) study [16], yielding a fragmentomics-based CSF-likeness score that discriminates between patients with favorable and unfavorable failure-free survival (Figures 4–6).

**Figure 2:**
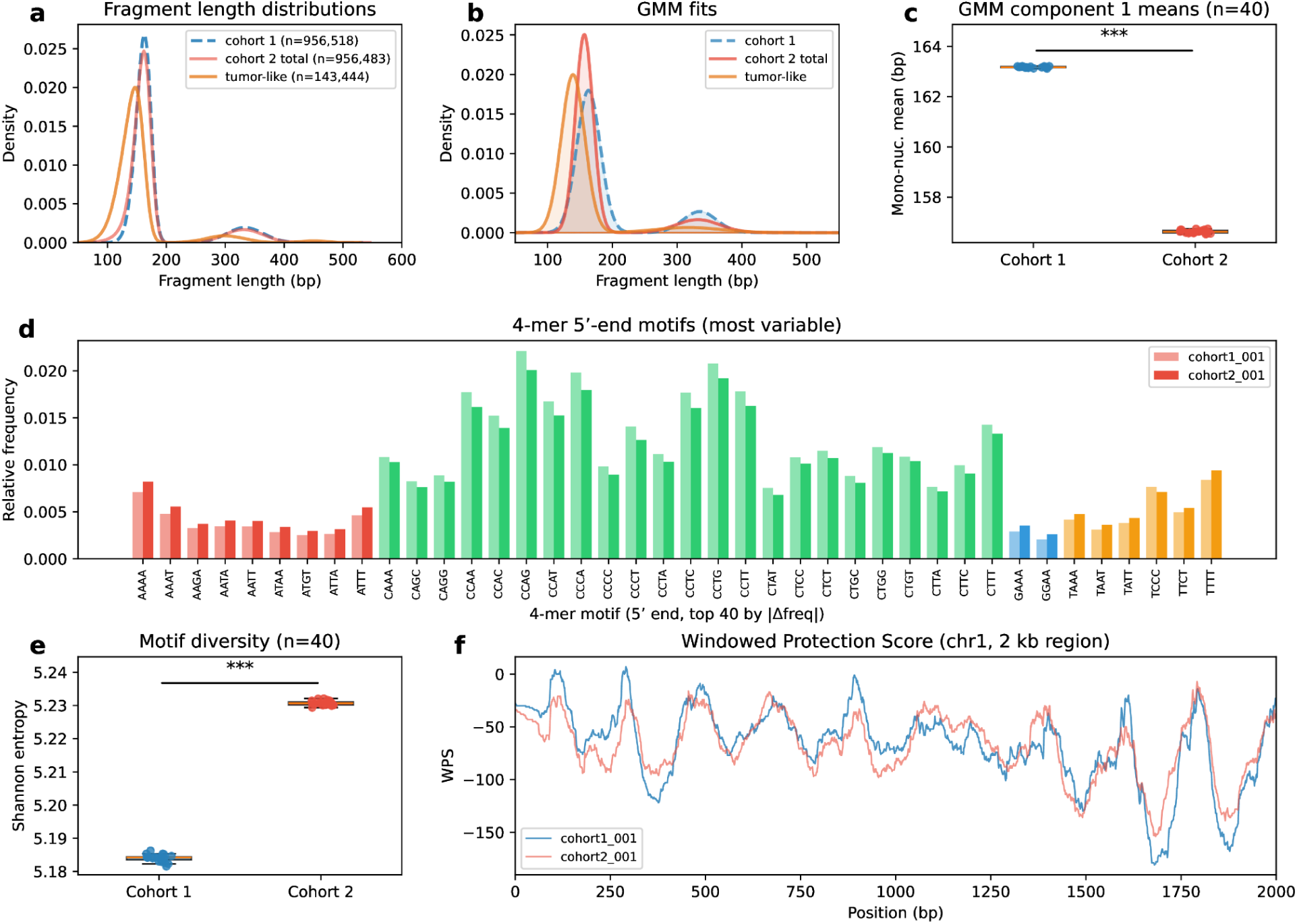
Per-sample analysis of two simulated cohorts of 20 samples each (cohort 1 is healthy-like, cohort 2 has an added 15% tumor-like admixture). (a) Fragment length kernel density estimates for one representative cohort 1 and one cohort 2 sample, the latter being stratified by mutation status. Fragment counts are given in the legend. (b) Gaussian mixture model fits of the same three fragment sets. Shaded areas indicate individual component contributions. (c) Fitted mononucleosomal component means across all 40 samples (n=20 per group). *** signifies p < 0.001, Mann-Whitney U test. (d) Relative frequencies of the 40 most variable 4-mer 5′ end motifs for one representative cohort 1 sample (translucent) and one cohort 2 sample (solid), sorted by absolute frequency difference. Bars are colored according to the first base of the respective motif. (e) Shannon entropy of 5′ end motif distributions across all 40 samples. *** signifies p < 0.001, Mann-Whitney U test. (f) Windowed protection score over a representative 2 kb genomic window on chromosome 1 for one cohort 1 sample (blue) and one cohort 2 sample (red).

### *Pyfraglib* accurately recovers simulated fragmentomic signatures

We first evaluated *pyfraglib* on simulated data, where the underlying fragmentomic differences are fully controlled and can effectively be used to validate functionality of the feature-extraction and cohort-level inference module. Therefore, we generated two cohorts of 20 synthetic cfDNA samples each. The first cohort is simulated to represent healthy plasma cfDNA as purely hematopoietic in origin, with DNASE1L3-dominant cleavage and a mononucleosomal fragment length distribution centered at 167 bp. Simulation hyperparameters for the second cohort are chosen to generate a mixture of 85% hematopoietic background that has parameters identical to the simulated normal samples, and 15% fragments characterized by strongly reduced DNASE1L3 activity, increased DFFB activity, and a shorter mean fragment length of 150 bp. These parameters recapitulate fragmentomics data found in tumor patients in the past [3, 28] and will thus be called tumor-like for the rest of this manuscript. Each sample contains roughly 1,000,000 simulated fragments, and fragments in the second cohort are explicitly labeled, providing ground-truth class membership for evaluation.

Figure 2 illustrates *pyfraglib’s* per-sample analysis capabilities applied to the simulated cohort. Fragment length distributions are visualized as kernel density estimates (Figure 2a). Three populations are shown: cohort 1 cfDNA (mononucleosomal peak at ∼167 bp), cohort 2 total (a mixture of fragments), and cohort 2 tumor-like fragments in isolation (centered near 150 bp). The shift in the tumor-like fragment length distribution relative to the rest of cohort 2 directly reflects the 15% admixture. GMM fitting (Figure 2b) decomposes each population into interpretable Gaussian components. Comparing the fitted mononucleosomal peak means across all 40 samples (Figure 2c) reveals the expected shift toward shorter fragments in cohort 2 relative to cohort 1 because the former contains tumor-like fragments (Mann-Whitney U test, p < 0.001). This result is consistent with the simulation design.

End motif profiles (Figure 2d) show the relative frequencies of the 40 most variable 4-mer 5′ end motifs across a representative cohort 1 and cohort 2 sample pair. Cohort 2 samples exhibit depletion of C-starting motifs attributable to the reduced DNASE1L3 and elevated DFFB activity of the tumor-like component. Shannon entropy of 5′ end motifs across all 40 samples (Figure 2e) confirms this at cohort scale (Mann-Whitney U test, p < 0.001), with cohort 2 samples showing lower end motif diversity as expected by construction from the altered nuclease balance.

The windowed protection score (Figure 2f) displays nucleosome-phased fragment coverage over a representative 2,000 bp genomic window. The signal shows regularly spaced peaks at approximately 200 bp intervals in both cohorts, consistent with nucleosome positioning. The similar patterns in cohort 1 and 2 samples reflect the dominant hematopoietic background (85%) shared by both groups.

Figure 3 demonstrates *pyfraglib’s* cohort-level analysis capabilities on both simulated cohorts. Differential end motif analysis (Figure 3a) compares all 512 4-mer end motifs (256 5′ and 256 3′) between the 20 cohort 1 and 20 cohort 2 samples using Wilcoxon rank-sum tests with Benjamini-Hochberg FDR correction. A total of 257 motifs are significantly differentially abundant (FDR < 0.05), representing approximately half of all tested motifs. This high proportion reflects the expected systematic and genome-wide nature of the simulated nuclease activity difference. The majority of significant motifs are C-starting sequences depleted in tumor-like relative to the other fragments, consistent with the reduced DNASE1L3-associated CC-cleavage activity in the tumor-like component.

**Figure 3:**
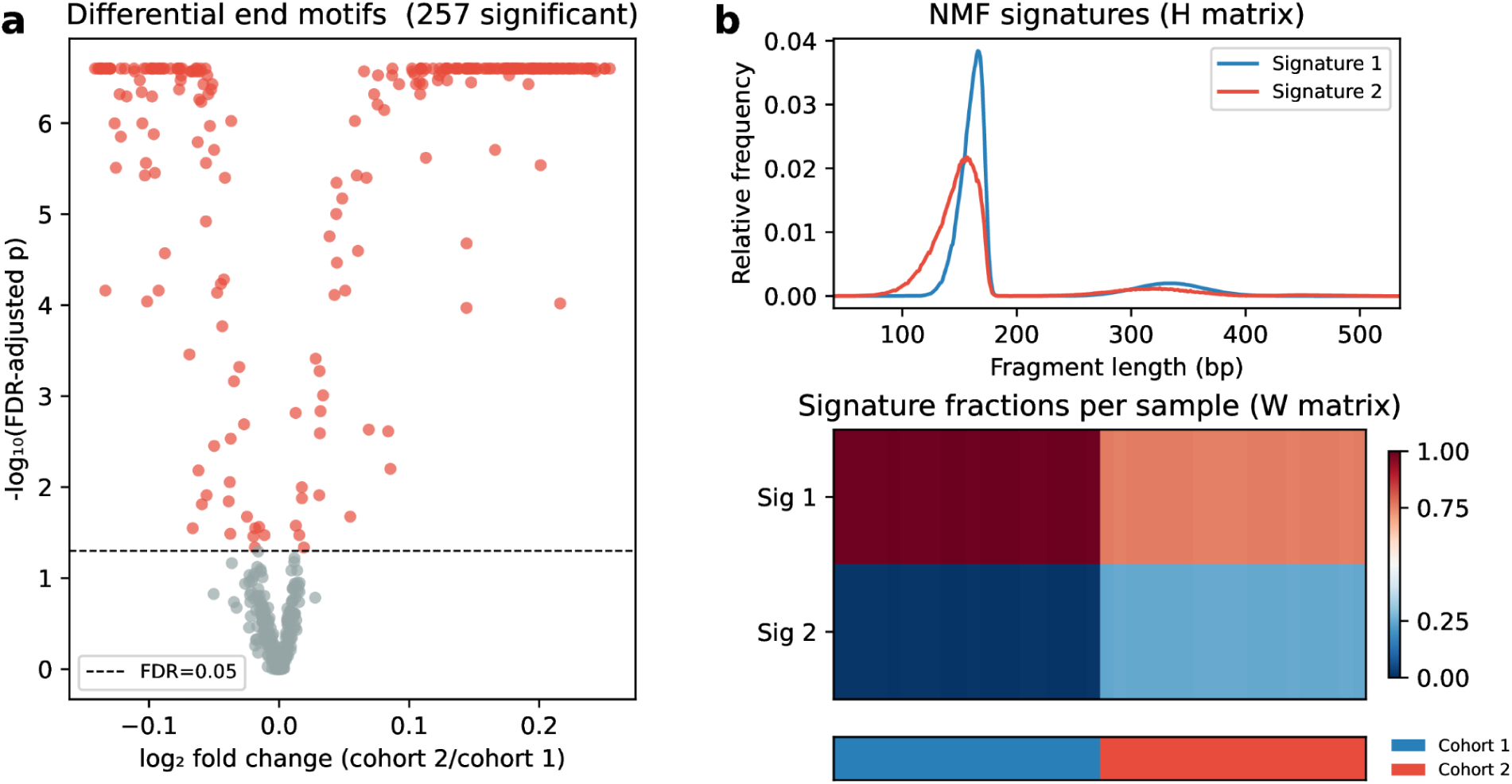
Cohort-level analysis of all 40 simulated samples (cohort 1 is healthy-like, cohort 2 has an added 15% tumor-like admixture). (a) Volcano plot of differential 4-mer end motif abundance between cohort 1 and cohort 2 (n=20 per group). x-axis: log₂ fold change (cohort 2/cohort 1), y-axis: −log₁₀(FDR-adjusted p-value, Benjamini-Hochberg). Motifs significant at FDR < 0.05 are shown in red, dashed line marks the significance threshold. (b) NMF decomposition of fragment length distributions into two signatures. Top: H matrix showing the two NMF signature profiles over fragment length. Bottom: W matrix heatmap of per-sample signature fractions (columns = samples, rows = signatures). Values represent the H-weighted fractional contribution of each signature. The annotation strip below indicates sample group (blue = cohort 1, red = cohort 2).

NMF decomposition of fragment length distributions across all 40 samples into two signatures (Figure 3b) yields components corresponding to background (Signature 1, mononucleosomal peak at ∼167 bp) and tumor-like (Signature 2, shifted peak at ∼150 bp with reduced periodicity) fragmentation patterns. The sample loading matrix stratifies samples consistently by cohort: all 20 cohort 1 samples load predominantly on Signature 1 while all 20 cohort 2 samples show elevated Signature 2 contribution. It is noteworthy that the recovered mean Signature 2 fraction for cohort 2 (∼0.25) exceeds the simulated 15% admixture. This overestimation is a known consequence of NMF when component signatures overlap substantially [29].

### *Pyfraglib* discovers fragmentomics signals in a CNSL cohort

To assess *pyfraglib’s* extraction pipeline on real clinical data, we exemplarily applied it to 88 independent cfDNA samples from healthy volunteers (n=10, CTRL) and CNSL patients (n=71 baseline (BL) plasma samples and n=7 cerebrospinal fluid (CSF) samples sequenced using a targeted gene panel [16] (Figure 4).

**Figure 4:**
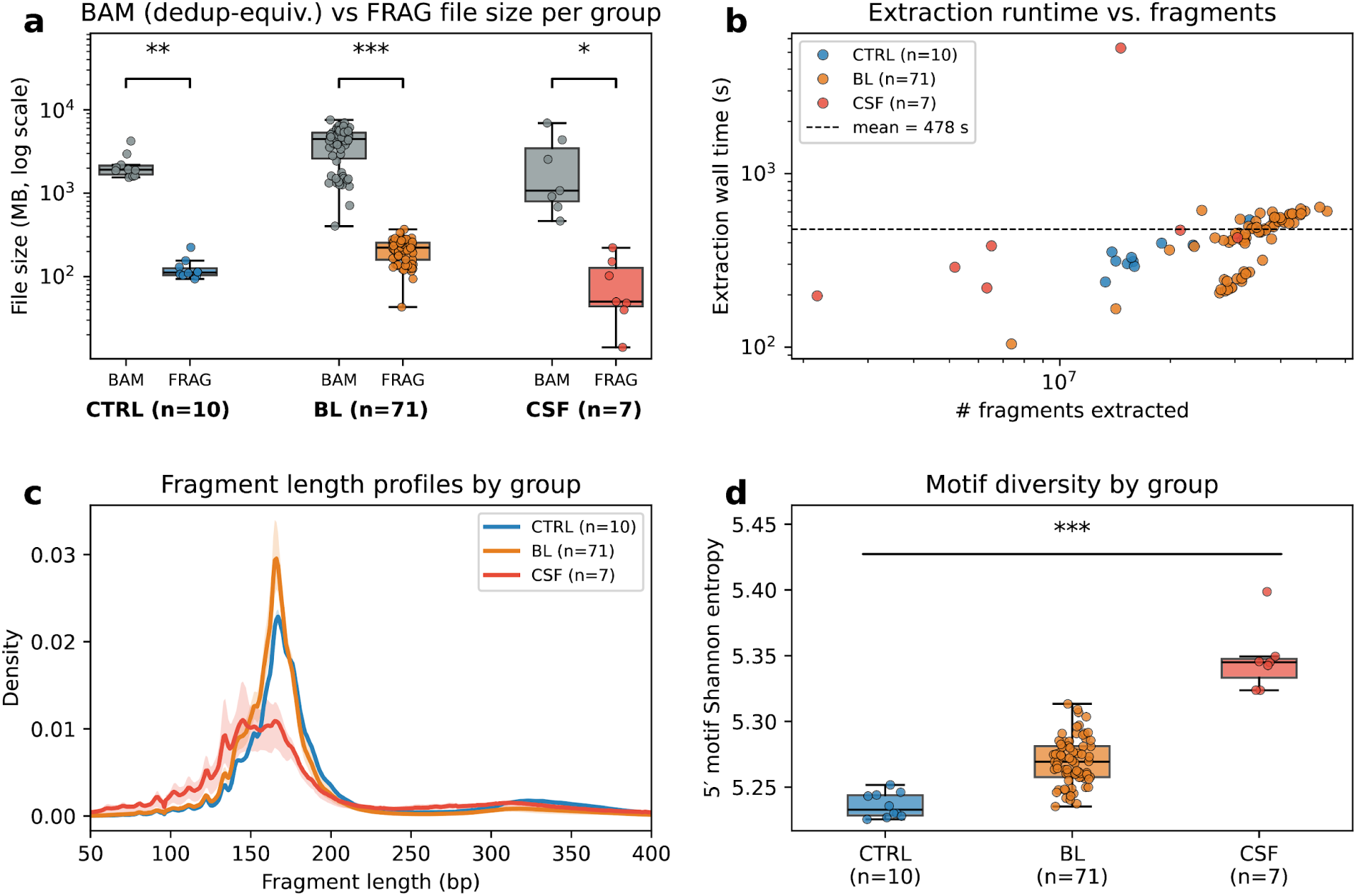
Real-data validation on ten healthy donor (CTRL) plasma cfDNA samples, 71 baseline (BL) samples, and seven cerebrospinal fluid (CSF) samples. (a) Duplicate-corrected BAM file sizes and FRAG file sizes per group, shown as boxplots over individual samples. Comparison was performed using paired Wilcoxon signed-rank tests. (b) Extraction wall time versus number of fragments per sample, processed on a 32-core HPC node. The dashed line indicates the cohort mean (478 seconds). (c) Per-group mean fragment length density with shaded interquartile range for CTRL (blue), BL (orange), and CSF samples (red). (d) Shannon entropy of 5′ end motif distributions per group. * signifies p < 0.05, ** signifies p < 0.01, *** signifies p < 0.001, Kruskal-Wallis test.

The compression of BAM into FRAG files led to a significantly reduced file size for all sample groups with a median compression ratio of 19.7 (Figure 4a), reflecting the intrinsic compactness of the FRAG format, which retains only information required for downstream fragmentomics analysis rather than raw sequencing reads and quality scores. Mean extraction time was 478 seconds (approximately 8 minutes) using 32 HPC cores. Additionally, it can be observed that runtime scales approximately linearly with BAM file size as indicated by the number of extracted fragments per second (Figure 4b). Fragment length distributions (Figure 4c) averaged across CTRL and BL samples show the characteristic plasma cfDNA profile observed in other studies [4], with empirical modes at approximately 167 bp and 320 bp which represent the expected mono- and dinucleosomal peaks. In contrast, no sharp mononucleosomal peak is observed when pooling CSF samples. On the contrary, the density estimate presents a plateau of most commonly observed fragment lengths between roughly 145 to 165 bp. The motif diversity of 5’ motifs was quantified by Shannon entropy (Figure 4d). While the entropy was lowest in the CTRL group, it slightly increased for the BL samples and even more so for the CSF group. The overall low inter-sample variance confirms the reproducibility of end motif extraction across independently processed samples.

To test *pyfraglib*’s clinical utility, we evaluated the cohort-level analysis workflow implemented in *pyfraglib* using clinical CNSL samples and healthy controls. For unsupervised comparison of the fragment lengths distributions (50-400 bp), NMF was applied to all CTRL and CSF samples (Figure 5a). The two identified signatures, S1, characterized by a CSF-enriched profile with a mononucleosomal peak at approximately 145 bp, and S2 with a CTRL-dominated peak at 167 bp, were then projected onto the BL samples. CSF samples showed heterogeneous loading patterns with an overall tendency toward S1, whereas CTRL and most BL samples predominantly loaded on S2. Notably, a subset of BL samples exhibited more balanced contributions from both signatures.

**Figure 5:**
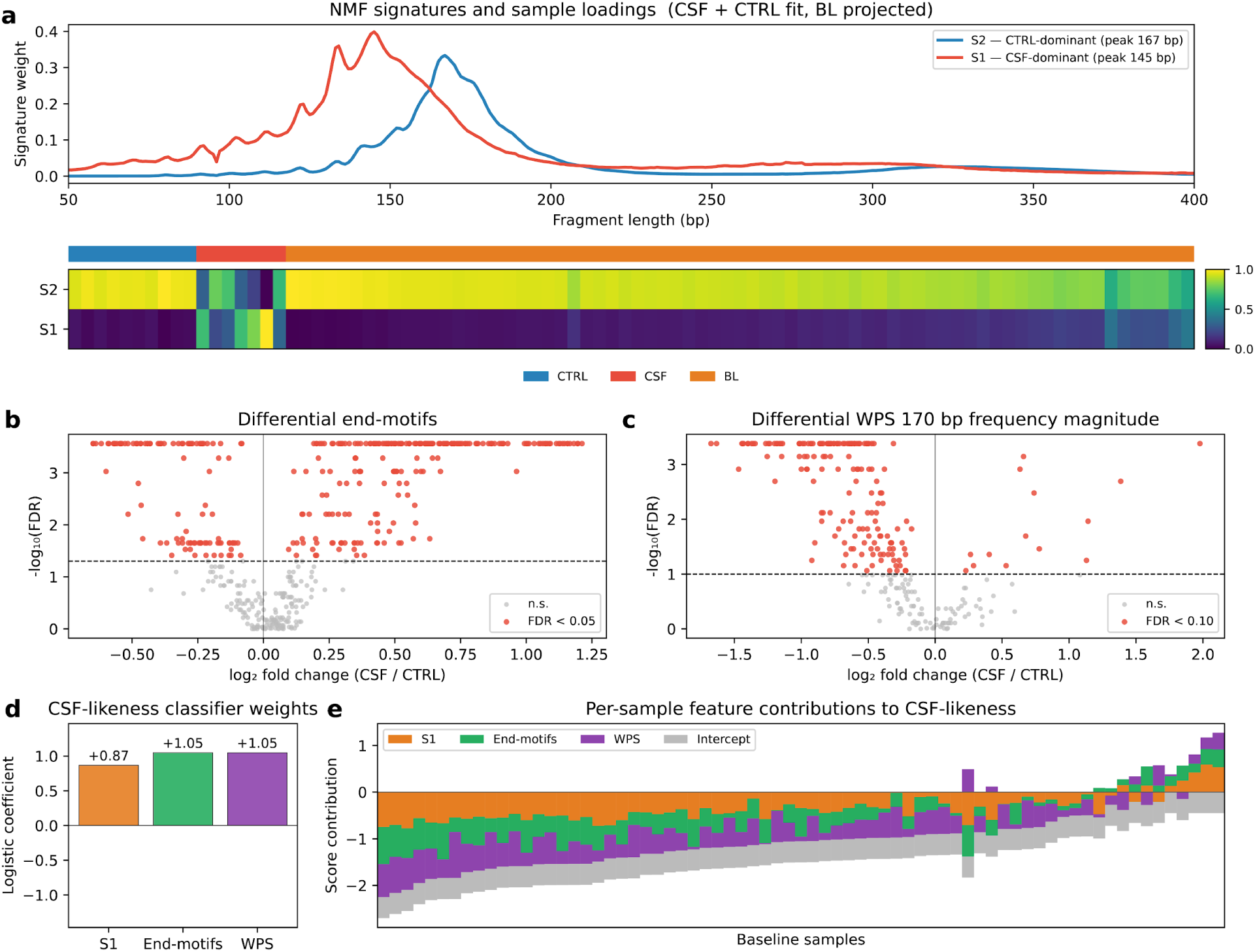
Differential analysis of fragmentomics features in CNSL patients. (a) Top: two NMF signatures extracted from the fragment length profiles of ten CTRL and seven CSF samples. Bottom: heatmap of per-sample signature loadings for CTRL, CSF, and the 71 BL samples. The latter are projected onto the fixed signatures. Group annotation strip below. (b) Volcano plot of differential 4-mer end motif abundance between CSF and CTRL samples (n=7 and n=10, respectively). x-axis: log₂ fold change (CSF/CTRL), y-axis: −log₁₀(FDR-adjusted p-value, Benjamini-Hochberg). Motifs significant at FDR < 0.05 are shown in red, a dashed line marks the significance threshold. (c) Volcano plot of differential WPS (per-region power at the ∼170 bp periodicity) between CSF and control samples (n=7 and n=10, respectively). x-axis: log₂ fold change (CSF/CTRL), y-axis: −log₁₀(FDR-adjusted p-value, Benjamini-Hochberg). Regions significant at FDR < 0.10 are shown in red, a dashed line marks the significance threshold. (d) Logistic-regression coefficients of the CSF-versus-CTRL classifier for its three input features: NMF signature 1 (orange), the end-motif score (green), the WPS score (purple). (e) Per-sample CSF-likeness scores for the 71 BL samples, shown as stacked contributions from the three input features and the classifier intercept.

Next we considered end-motifs. At a Wilcoxon rank-sum FDR of 0.05, the differential analysis of end-motifs comparing CSF to CTRL samples revealed a higher number of more abundant motifs in CSF samples (225 motifs) as opposed to CTRL samples (107 motifs, Figure 5b). To summarize each sample’s end motif profile, a per-sample score was defined as the signed sum of z-scored 4-mer frequencies across the significant end motifs only (Supp. Figure 3a).

Lastly, analysis of windowed protection scores (WPS) across the 284 genomic regions (the union of all panel-covered genomic intervals that pertain to a distinct gene [16]) revealed a striking asymmetry between CSF and CTRL samples (Figure 5c). Per-region power spectrum ratio (see Methods) z-scores are weighted by the sign of the region’s CSF-vs-CTRL log-fold-change and summed over regions significant at FDR < 0.10. Of the 178 regions significantly different between groups with respect to the spectral power feature (FDR < 0.10), the vast majority were elevated in CTRL samples (164 regions). Analogously to the end motifs, a per-sample WPS score was defined as the sum of z-scored feature values across the significant regions (Supp. Figure 3b).

To derive a unified CSF-likeness score, the three z-scaled cohort-level features (CSF-associated length signature loading, end-motif score, and WPS score) were combined with a logistic classifier with L2 regularisation, trained to discriminate CSF from CTRL. The resulting classifier weights were all positive and similar in magnitude (range 0.87 - 1.05, Figure 5d). Subsequently applying the fixed classifier to the baseline samples then yielded a CSF-likeness score for each BL sample (Figure 5e). Because weights were identical across samples, the per-sample differences reflect each sample’s underlying feature values. Contributions varied considerably between baseline samples, with no single feature dominating. An overview of the three features and the resulting score for every sample is provided in Supp. Figure 2.

To illustrate a potential clinical application of such a classifier built using *pyfraglib’s* cohort-level analyses, the distribution of CSF-likeness scores across a sub-cohort of 66 baseline samples was visualized with respect to failure-free survival (FFS) status of each patient as defined by Heger et al. [16] (Figure 6a). For this analysis, 5 samples that constituted repeated measurements from the same patients were removed.

**Figure 6:**
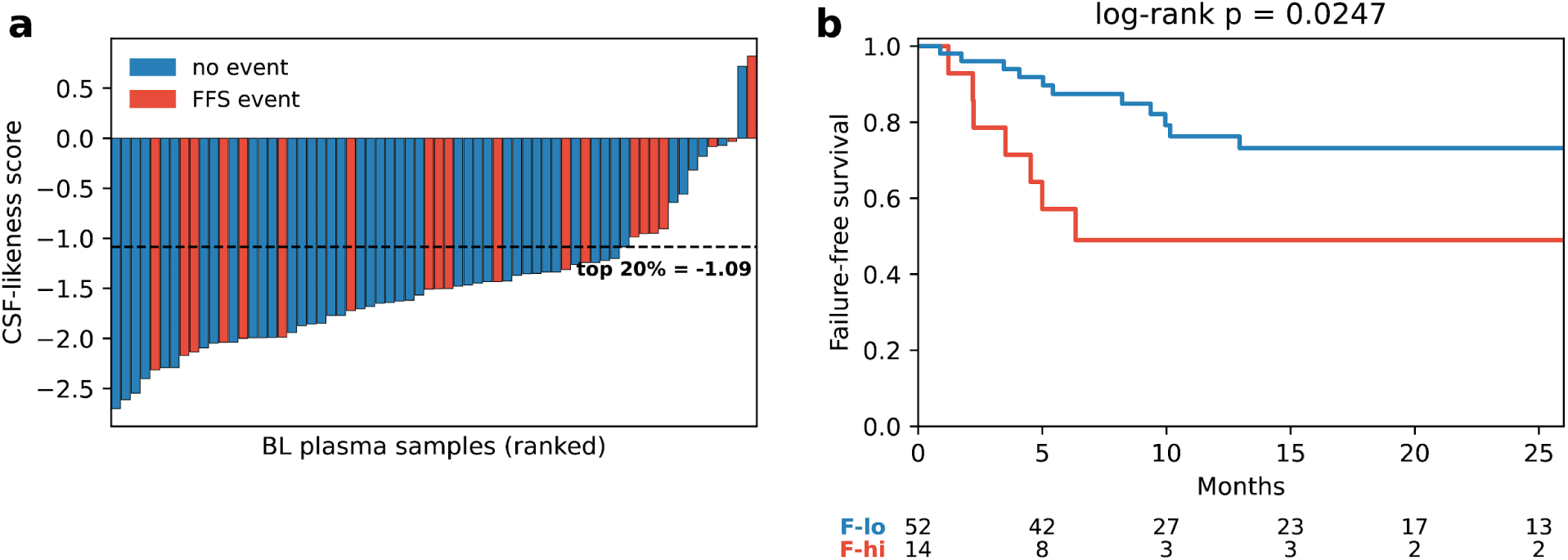
Association of CSF-likeness classifier with failure-free survival in CNSL patients. (a) Per-sample CSF-likeness score for 66 BL samples, ranked in ascending order and color-coded according to failure-free survival (FFS) status (red = event, blue = no event). The dashed line marks the exploratory 20% cutoff for stratification used in panel (b). (b) Kaplan-Meier of FFS comparing the top-20% CSF-likeness-high (F-hi, n = 14, red) with the remaining CSF-likeness-low samples (F-lo, n = 52, blue), log-rank p = 0.0247.

Then, we stratified the 66 individuals into a CSF-likeness-high (F-high) and CSF-likeness-low (F-low) group using an exploratory 20%/80% split. The Kaplan-Meier plot in Figure 6b confirms that in this cohort, a CSF-likeness score above the 80th percentile is significantly associated with an unfavorable FFS (log-rank (Davidson-Pilon, 2019) p = 0.0247). Exploratory testing for significant associations with clinical characteristics revealed high CSF-likeness to be linked to female sex, high ctDNA and cfDNA content of respective samples, and poor performance status at relapse (Mann-Whitney U tests for continuous and a χ² test for categorical variables, with Benjamini-Hochberg FDR, Supp. Figure 4).

Together, these results demonstrate that *pyfraglib* provides a unified framework for fragmentomics analysis across both controlled and clinical settings at single sample as well as cohort level. Validation on simulated cohorts demonstrates the accuracy of its analytical methods under known ground truth, whereas application to a clinical CNSL cohort showcases its ability to extract biologically and clinically relevant fragmentomic signatures from patient data. These findings illustrate the utility of *pyfraglib* for method development, biological discovery, and translational cfDNA research.

## Discussion

In this paper, we present *pyfraglib*, an integrated Python platform for cfDNA fragmentomics analysis combining fragment extraction into an efficient intermediate file format, statistical feature modeling, cohort-level comparative analysis, and *in silico* simulation. Relative to existing tools, *pyfraglib* is the first single package to combine WPS calculation, GMM fitting of fragment length distributions, NMF decomposition, as well as end motif diversity quantification and differential testing on cohort-level within a consistently engineered Python framework.

The simulation module extends *pyfraglib* beyond a feature extraction toolkit into a framework for fragmentomics method development and validation. By probabilistic modeling of nuclease cleavage of a template DNA and drawing fragment lengths from a controlled distribution, it enables generation of datasets with known ground truths for software testing, algorithm prototyping, and study design prior to committing to patient sample collection. The demonstration presented here - a cross-sectional cohort of 40 simulated samples with a 15% tumor-like admixture - confirms for the algorithms already implemented in *pyfraglib* that they recover the differences introduced by the simulation. Such simulation-based benchmarking directly extends to the development of novel algorithms which is facilitated by the accessible, well documented and extensively tested API of *pyfraglib*.

As an important caveat, exact mixture fraction recovery through NMF requires each signature to dominate exclusively in at least one feature, which the largely shared mononucleosomal peak structure of the two tissue types violates [30]. Thus, the simulation-based demonstration does not recover the simulated fraction of 15% exactly. To prove that this is an identifiability issue inherent to NMF and not caused by an incorrect implementation, *pyfraglib* includes unit tests that show exact mixture fraction recovery for cases of perfectly separated signatures.

We furthermore emphasize that the simulation parameters are biologically motivated but not exhaustively calibrated. Users requiring parameters closely matched to a specific assay or cohort should optimize them against real data before using the simulation for power analyses or algorithm comparison. In this study, we aim to verify that the implemented analyses behave as expected under controlled conditions. This also ensures that *pyfraglib*’s commands are fully reproducible without access to patient-derived or controlled-access datasets.

Validation on 88 real-world cfDNA samples (Figures 4-6) demonstrated that the extraction pipeline operates robustly on clinical data, generating biologically plausible fragment length distributions and end motif profiles consistent with the published cfDNA literature. The median BAM-to-FRAG compression ratio of roughly 20x highlights the practical storage and data-handling advantages of the FRAG file format for large-scale cfDNA cohorts.

As a proof of principle, cohort-level differential analysis of the extracted fragmentation features further demonstrated the potential of *pyfraglib*-derived metrics to capture clinically relevant signals and provide prognostic value. Notably, in the regularized CSF-likeness score all features received comparable weights, thus no single feature dominated the classifier. This finding underscores the value of a joint analysis of multiple fragmentomics features as done by *pyfraglib*. Importantly, the presented CSF-likeness score is meant to be an illustrative example only. It is resting on a single, modestly sized cohort and has not been validated independently. Additional studies would be required to establish clinical meaningfulness.

Several limitations of the current implementation should be noted. First, *pyfraglib* does not perform GC bias correction, which can confound fragment length and WPS estimates particularly at GC-rich genomic regions. Complementary tools such as GCparagon [31] or GCfix [32] can be applied upstream. Second, methylation-based fragmentomics features are outside the current scope. Lastly, while the test suite covers core functionality, coverage of edge cases and rare biological scenarios will grow as the tool matures. Addressing these limitations is part of our future development roadmap.

## Conclusions

*pyfraglib* provides an integrated, open-source Python platform for cfDNA fragmentomics analysis that unifies fragment extraction, statistical feature modeling, cohort-level comparative analysis, and *in silico* simulation within a single consistently engineered package. By making all of these capabilities available through a common API, CLI, and Nextflow pipeline, *pyfraglib* lowers the barrier to entry for groups seeking to apply or develop fragmentomics methods and removes the need to integrate multiple incompatible tools. The *in silico* simulation module enables rigorous algorithm benchmarking and study design without patient-derived data. Validation on real cfDNA samples confirms that the extraction pipeline produces results consistent with the published cfDNA literature and indicated translational potential. We anticipate that future development will extend *pyfraglib* with GC bias correction, methylation-based algorithms and more realistic simulation capabilities, further broadening its applicability in both research and clinical translation contexts.

## Availability and requirements

Project name: pyfraglib

Project home page: https://github.com/schwarzlab-ccb/pyfraglib

Operating system(s): Platform independent

Programming language: Python

Other requirements: none

License: GNU General Public License v3

Any restrictions to use by non-academics: Licence needed

## List of abbreviations

API: application programming interface
BAM: Binary Alignment Map (file format)
BED: Browser Extensible Data (file format)
BL: baseline (plasma sample)
bp: base pair(s)
cfDNA: cell-free DNA
CLI: command-line interface
CNSL: central nervous system lymphoma
CSF: cerebrospinal fluid
ctDNA: circulating tumor DNA
CTRL: (healthy-donor) control
DFFB: DNA fragmentation factor subunit beta
DNA: deoxyribonucleic acid
DNASE1: deoxyribonuclease 1
DNASE1L3: deoxyribonuclease 1-like 3
FASTA: text-based nucleotide sequence format
FDR: false discovery rate
FFS: failure-free survival
F-hi / F-lo: fragmentomics-high / fragmentomics-low subgroup
FRAG: *pyfraglib* fragment file format
GC: guanine–cytosine
GMM: Gaussian mixture model
HPC: high-performance computing
MOP-C: molecular prognostic index for CNS lymphoma [16]
NMF: non-negative matrix factorization
SD: standard deviation
VCF: Variant Call Format
WPS: windowed protection score

## Declarations

### Ethics Approval and Consent to Participate

The 88 cfDNA samples used for real-data validation were obtained from an independent study [16]. Ethics approval was granted in the context of the respective study. All participants provided written informed consent prior to sample collection.

### Consent for publication

Not applicable.

### Availability of data and materials

An archived version of the scripts used to generate the figures in this manuscript is deposited at https://doi.org/10.5281/zenodo.21720982. The simulated datasets used in the *in silico* demonstration can be fully reproduced using the configuration files provided in the *config* directory of the *pyfraglib* Github repository. The clinical samples and metadata from the Heger et al. study [16] are not publicly available due to patient privacy restrictions but may be requested from the corresponding author subject to data access agreement.

### Authors’ Contributions

DS: Conceptualization, Software architecture and implementation, Formal analysis, Visualization, Writing – original draft, Funding acquisition. LKG: Testing of Software, Conceptualization, Writing - review & editing. JS: Testing of Software, Conceptualization, Writing - review & editing. SB: Resources, including healthy donor samples, Conceptualization, Writing - review & editing, Funding acquisition. JMH: Resources, including healthy donor samples, Writing – review & editing, Funding acquisition. RFS: Conceptualization, Supervision, Funding acquisition, Writing – review & editing. All authors read and approved the final manuscript.

## Acknowledgements

We gratefully acknowledge Adam Streck for his valuable comments regarding the implementation of *pyfraglib* and the present manuscript. We thank the Regional Computing Center of the University of Cologne (RRZK) for providing computing time on the High Performance Computing (HPC) system RAMSES as well as support.

The large language model Claude Sonnet 4.6 (Anthropic) was used for proof-reading of this manuscript. The large language model GPT-5.2 (OpenAI) was used to generate the *pyfraglib* logo.

## Supplementary Material

**Supplementary Figure 1:**
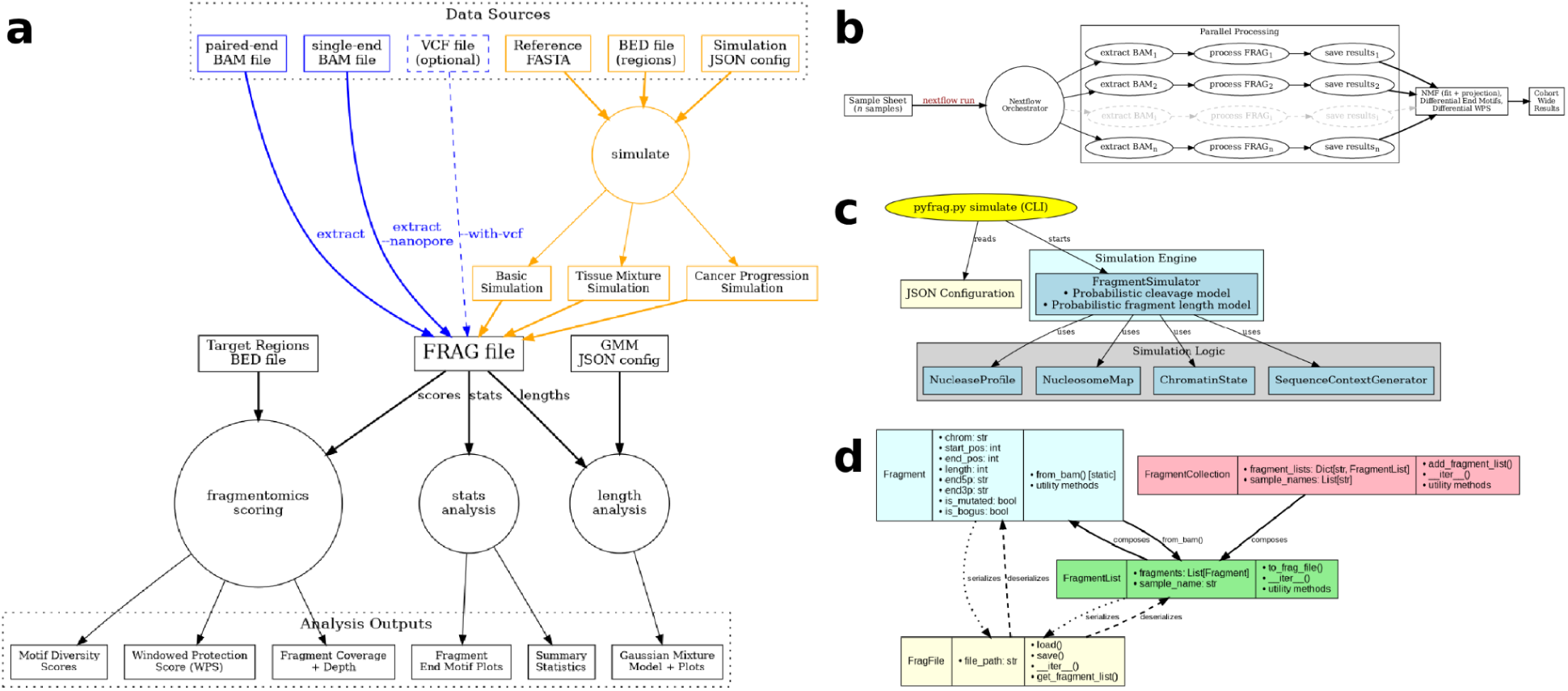
Architecture overview. (a) End-to-end workflow diagram showing supported input sources (paired-end and single-end BAM files, optional VCF, reference FASTA, BED regions, simulation JSON configuration), the central FRAG file format, and the three analysis branches (fragmentomics scoring, stats analysis, length analysis) together with their outputs. (b) Graphical overview of Nextflow pipeline for HPC-scale cohort processing: a sample sheet is passed to the Nextflow orchestrator, which parallelizes extract, process, and save-results steps across n samples before aggregating cohort-wide NMF and differential end motif results. (c) Simulation architecture: the **simulate** CLI subcommand reads a JSON configuration and drives the FragmentSimulator (probabilistic cleavage and fragment length models), which relies on NucleaseProfile, NucleosomeMap, ChromatinState, and SequenceContextGenerator. (d) Core Python API class diagram illustrating the Fragment, FragmentList, FragmentCollection, and FragFile classes, their fields, key methods, and compositional relationships.

**Supplementary Figure 2:**
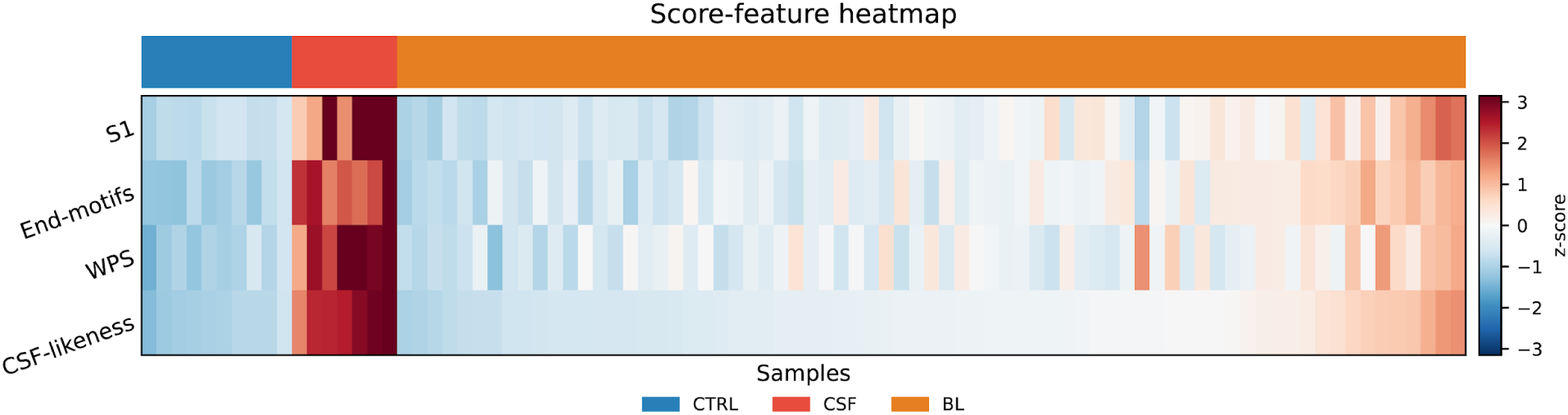
Per-sample heatmap of the three CSF-likeness score features and the resulting score across all 88 samples. Rows show the CSF-signature loading (S1), the end-motif score, and the WPS score, together with the final CSF-likeness score (logistic-regression decision value). Each row is z-scored across samples (color = z-score). Columns are the 88 samples, sorted within groups (CTRL, CSF, BL) by CSF-likeness score, with a group-annotation strip above.

**Supplementary Figure 3:**
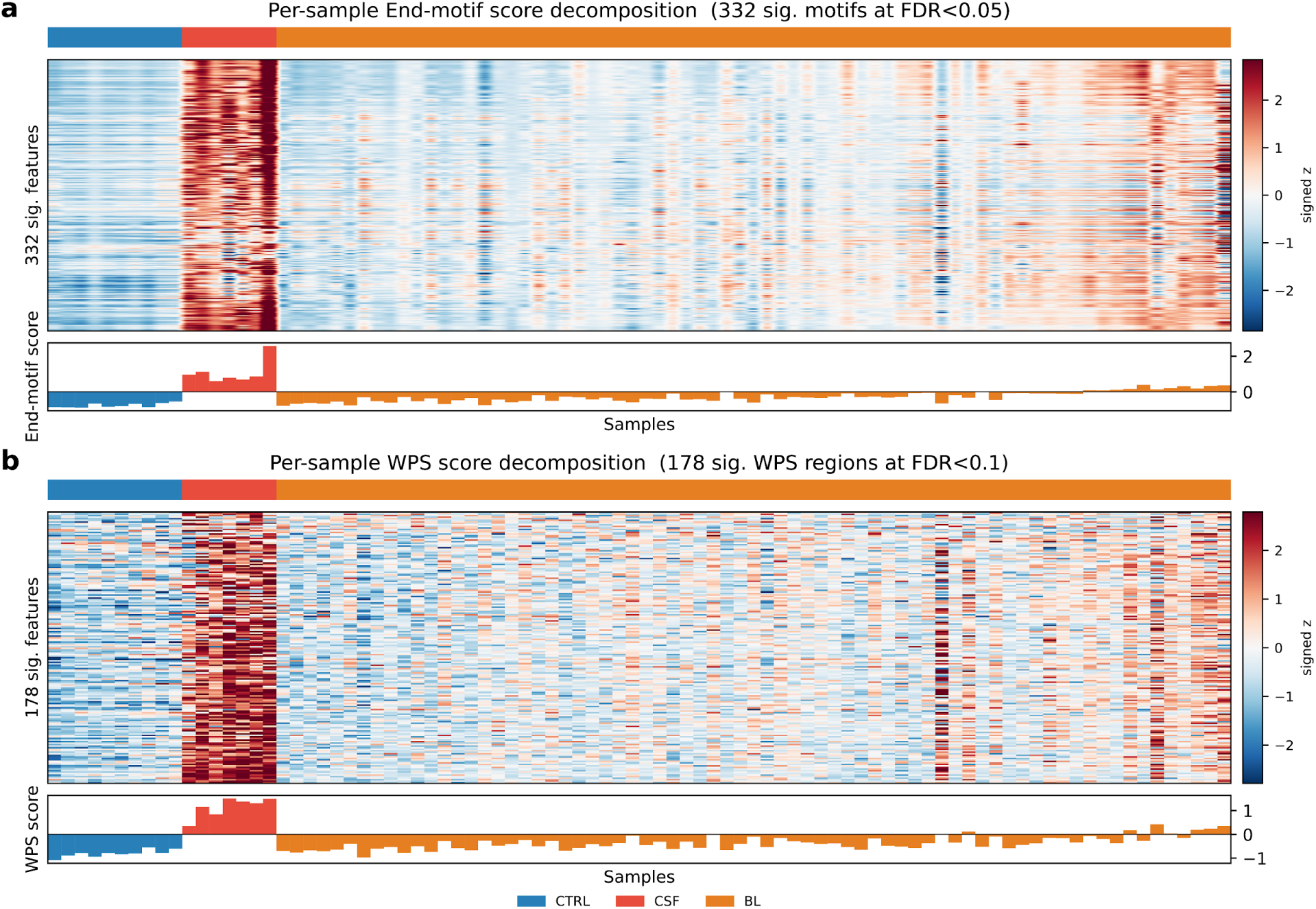
Per-sample construction of the end-motif and WPS scores. (a) End-motif score decomposition across the 332 end-motif features significantly different between CSF and CTRL (FDR < 0.05). (b) WPS score decomposition across the 178 regions significantly different between CSF and CTRL (FDR < 0.10). In each panel, heatmap rows are the significant features, sorted by log₂ fold change (most CSF-enriched at top). Each cell is the signed z-score contribution to the score [sign(log₂ fold change) × z of the per-sample feature value], with red indicating CSF-like (positive) and blue non-CSF-like (negative) contributions. Columns are 88 samples, sorted within groups (CTRL, CSF, BL) by CSF-likeness score, with a group-annotation strip above. The bar plot beneath each heatmap is the per-sample score (the column sum of the heatmap) used in the CSF-likeness classifier.

**Supplementary Figure 4:**
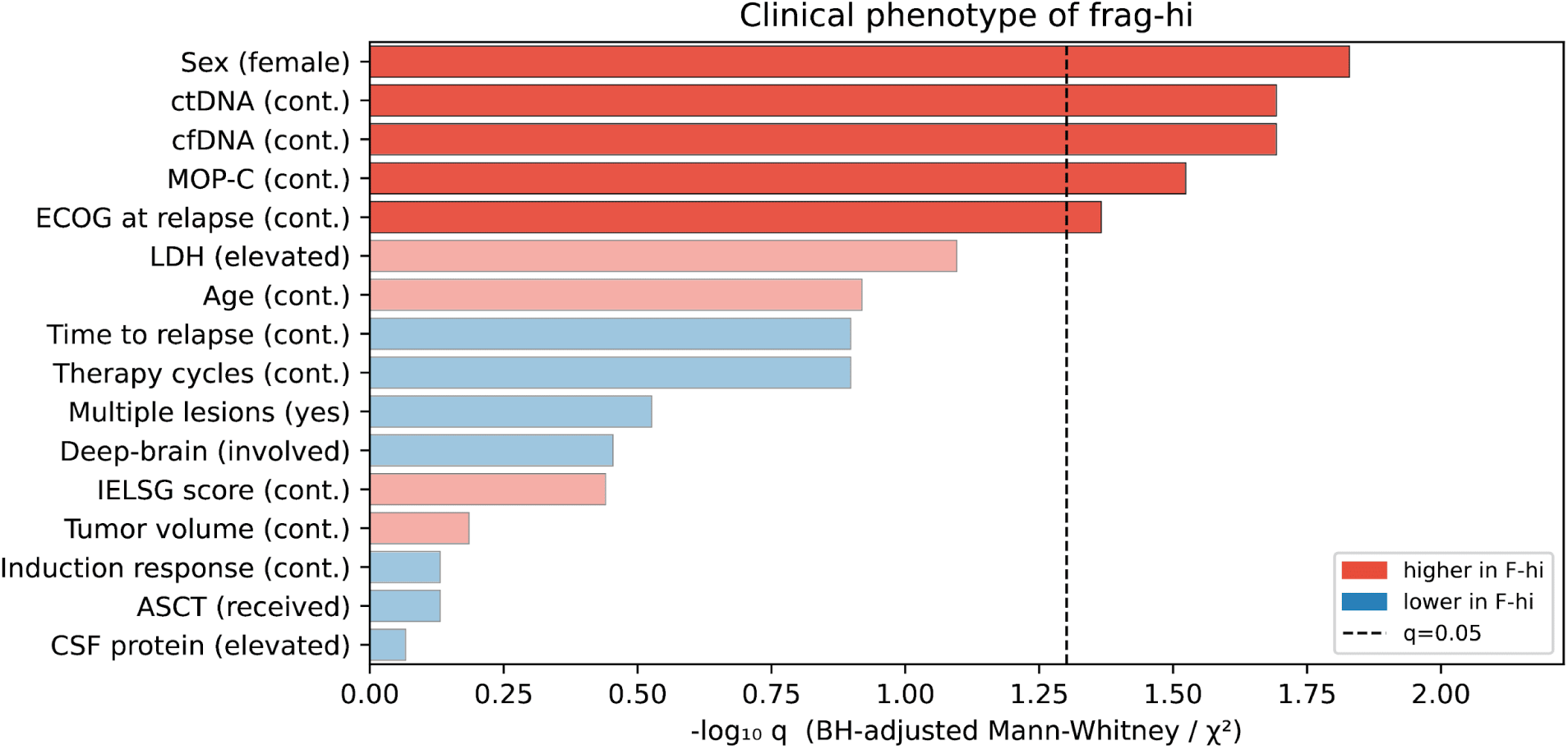
Clinical phenotype of the CSF-likeness-high baseline subgroup. Per-variable comparison of the top-20% CSF-likeness baseline samples (F-hi, n = 14) versus the remaining baseline samples (F-lo, n = 52). Bars show −log₁₀ of the Benjamini-Hochberg-adjusted p-value or each of 16 clinical variables, tested by Mann-Whitney U (continuous variables) or χ² (categorical variables). The dashed vertical line marks q = 0.05. Bars are colored by direction and significant variables (q < 0.05) are drawn at full opacity. For categorical variables, the reference class is given in the y-axis label.

## Bibliography

1. Lo YMD, Han DSC, Jiang P, Chiu RWK. Epigenetics, fragmentomics, and topology of cell-free DNA in liquid biopsies. Science. 2021;372:eaaw3616. 10.1126/science.aaw3616.

2. Wan JCM, Massie C, Garcia-Corbacho J, Mouliere F, Brenton JD, Caldas C, et al. Liquid biopsies come of age: towards implementation of circulating tumour DNA. Nat Rev Cancer. 2017;17:223–38. 10.1038/nrc.2017.7.

3. Mouliere F, Chandrananda D, Piskorz AM, Moore EK, Morris J, Ahlborn LB, et al. Enhanced detection of circulating tumor DNA by fragment size analysis. Sci Transl Med. 2018;10:eaat4921. 10.1126/scitranslmed.aat4921.

4. Snyder MW, Kircher M, Hill AJ, Daza RM, Shendure J. Cell-free DNA Comprises an In Vivo Nucleosome Footprint that Informs Its Tissues-Of-Origin. Cell. 2016;164:57–68. 10.1016/j.cell.2015.11.050.

5. Ulz P, Thallinger GG, Auer M, Graf R, Kashofer K, Jahn SW, et al. Inferring expressed genes by whole-genome sequencing of plasma DNA. Nat Genet. 2016;48:1273–8. 10.1038/ng.3648.

6. Cristiano S, Leal A, Phallen J, Fiksel J, Adleff V, Bruhm DC, et al. Genome-wide cell-free DNA fragmentation in patients with cancer. Nature. 2019;570:385–9. 10.1038/s41586-019-1272-6.

7. Esfahani MS, Hamilton EG, Mehrmohamadi M, Nabet BY, Alig SK, King DA, et al. Inferring gene expression from cell-free DNA fragmentation profiles. Nat Biotechnol. 2022;40:585–97. 10.1038/s41587-022-01222-4.

8. Foda ZH, Annapragada AV, Boyapati K, Bruhm DC, Vulpescu NA, Medina JE, et al. Detecting Liver Cancer Using Cell-Free DNA Fragmentomes. Cancer Discov. 2023;13:616–31. 10.1158/2159-8290.CD-22-0659.

9. Mathios D, Johansen JS, Cristiano S, Medina JE, Phallen J, Larsen KR, et al. Detection and characterization of lung cancer using cell-free DNA fragmentomes. Nat Commun. 2021;12:5060. 10.1038/s41467-021-24994-w.

10. Wang H, Mennea PD, Chan YKE, Cheng Z, Neofytou MC, Surani AA, et al. A standardized framework for robust fragmentomic feature extraction from cell-free DNA sequencing data. Genome Biol. 2025;26:141. 10.1186/s13059-025-03607-5.

11. Li JW, Bandaru R, Baliga K, Liu Y. FinaleToolkit: accelerating cell-free DNA fragmentation analysis with a high-speed computational toolkit. Bioinforma Adv. 2025;5:vbaf236. 10.1093/bioadv/vbaf236.

12. Doebley A-L, Ko M, Liao H, Cruikshank AE, Santos K, Kikawa C, et al. A framework for clinical cancer subtyping from nucleosome profiling of cell-free DNA. Nat Commun. 2022;13:7475. 10.1038/s41467-022-35076-w.

13. Moldovan N, van der Pol Y, van den Ende T, Boers D, Verkuijlen S, Creemers A, et al. Multi-modal cell-free DNA genomic and fragmentomic patterns enhance cancer survival and recurrence analysis. Cell Rep Med. 2024;5:101349. 10.1016/j.xcrm.2023.101349.

14. Lazzeri I, Spiegl BG, Hasenleithner SO, Speicher MR, Kircher M. LBFextract: Unveiling transcription factor dynamics from liquid biopsy data. Comput Struct Biotechnol J. 2024;23:3163–74. 10.1016/j.csbj.2024.08.007.

15. Weber LM, Saelens W, Cannoodt R, Soneson C, Hapfelmeier A, Gardner PP, et al. Essential guidelines for computational method benchmarking. Genome Biol. 2019;20:125. 10.1186/s13059-019-1738-8.

16. Heger J-M, Mattlener J, Schneider J, Gödel P, Sieg N, Ullrich F, et al. Entirely noninvasive outcome prediction in central nervous system lymphomas using circulating tumor DNA. Blood. 2024;143:522–34. 10.1182/blood.2023022020.

17. Dempster AP, Laird NM, Rubin DB. Maximum Likelihood from Incomplete Data Via the *EM* Algorithm. J R Stat Soc Ser B Stat Methodol. 1977;39:1–22. 10.1111/j.2517-6161.1977.tb01600.x.

18. Shannon CE. A Mathematical Theory of Communication. Bell Syst Tech J. 1948;27:379–423. 10.1002/j.1538-7305.1948.tb01338.x.

19. Simpson EH. Measurement of Diversity. Nature. 1949;163:688–688. 10.1038/163688a0.

20. Savitzky Abraham, Golay MJE. Smoothing and Differentiation of Data by Simplified Least Squares Procedures. Anal Chem. 1964;36:1627–39. 10.1021/ac60214a047.

21. Cooley JW, Tukey JW. An algorithm for the machine calculation of complex Fourier series. Math Comput. 1965;19:297–301. 10.1090/S0025-5718-1965-0178586-1.

22. Lee DD, Seung HS. Learning the parts of objects by non-negative matrix factorization. Nature. 1999;401:788–91. 10.1038/44565.

23. Renaud G, Nørgaard M, Lindberg J, Grönberg H, De Laere B, Jensen JB, et al. Unsupervised detection of fragment length signatures of circulating tumor DNA using non-negative matrix factorization. eLife. 2022;11:e71569. 10.7554/eLife.71569.

24. Mann HB, Whitney DR. On a Test of Whether one of Two Random Variables is Stochastically Larger than the Other. Ann Math Stat. 1947;18:50–60. 10.1214/aoms/1177730491.

25. Benjamini Y, Hochberg Y. Controlling the False Discovery Rate: A Practical and Powerful Approach to Multiple Testing. J R Stat Soc Ser B Stat Methodol. 1995;57:289–300. 10.1111/j.2517-6161.1995.tb02031.x.

26. Han DSC, Lo YMD. The Nexus of cfDNA and Nuclease Biology. Trends Genet. 2021;37:758–70. 10.1016/j.tig.2021.04.005.

27. Di Tommaso P, Chatzou M, Floden EW, Barja PP, Palumbo E, Notredame C. Nextflow enables reproducible computational workflows. Nat Biotechnol. 2017;35:316–9. 10.1038/nbt.3820.

28. Serpas L, Chan RWY, Jiang P, Ni M, Sun K, Rashidfarrokhi A, et al. Dnase1l3 deletion causes aberrations in length and end-motif frequencies in plasma DNA. Proc Natl Acad Sci U S A. 2019;116:641–9. 10.1073/pnas.1815031116.

29. Fu X, Huang K, Sidiropoulos ND. On Identifiability of Nonnegative Matrix Factorization. IEEE Signal Process Lett. 2018;25:328–32. 10.1109/LSP.2018.2789405.

30. Jin H, Gulhan DC, Geiger B, Ben-Isvy D, Geng D, Ljungström V, et al. Accurate and sensitive mutational signature analysis with MuSiCal. Nat Genet. 2024;56:541–52. 10.1038/s41588-024-01659-0.

31. Spiegl B, Kapidzic F, Röner S, Kircher M, Speicher MR. GCparagon: evaluating and correcting GC biases in cell-free DNA at the fragment level. NAR Genomics Bioinforma. 2023;5:lqad102. 10.1093/nargab/lqad102.

32. Rahman CR, Poh ZW, Skanderup AJ, Wong L. GCfix: a fast and accurate fragment length-specific method for correcting GC bias in cell-free DNA. Bioinformatics. 2025;41:btaf293. 10.1093/bioinformatics/btaf293.

